# Genetic Diversity and Phylogenetic Relationships of Cultivated and Wild *Urochloa* Species Conserved in the ILRI Forage Genebank

**DOI:** 10.64898/2025.12.20.695694

**Authors:** Meki S. Muktar, Shimu D. Lema, Alemayehu Teressa Negawo, Yilikal Assefa, Helen Nigussie, Hailu M. Tola, Chris S. Jones

## Abstract

Understanding the genetic diversity and evolutionary relationships within the genus *Urochloa* is essential for effective germplasm conservation and the development of improved forage cultivars. In this study, we assessed the genetic diversity, population structure, and phylogenetic relationships of 545 accessions representing 17 *Urochloa* species conserved in the ILRI Forage Genebank for the last four decades at its Zway field site in Ethiopia. Large numbers of genome-wide SNP and SilicoDArT markers were generated by the GBS based Diversity Arrays Technology (DArT)seq platform, providing valuable resources for genetic analysis in *Urochloa* spp. From these, we filtered a subset of high-quality SNP and SilicoDArT markers suitable for cross-species analysis, which was then used to characterize species differentiation and to detect admixture patterns across wild and cultivated species. Hierarchical clustering grouped all 545 accessions into two major groups, clearly separating the brizantha complex (*U. brizantha, U. decumbens,* and *U. ruziziensis*) from the rest of the species, which were primarily wild and less domesticated relatives. The analysis further identified four well-defined phylogenetic clades, largely consistent with previously reported evolutionary relationships within the genus. The three clades, primarily composed of wild species, exhibited clear genetic separation, high diversity, and strong geographic structuring with limited admixture, reflecting reproductive isolation (their apomictic nature) and differing ploidy levels that have driven local adaptation. In contrast, the brizantha complex exhibited extensive admixture and comparatively lower genetic variation, consistent with their historical polyploidization events, shared ancestry, and domestication (particularly in the key forage species *U. brizantha* and *U. decumbens*). Population structure analysis using the admixture model revealed multiple genetically distinct clusters and populations across clades, including species-specific clusters, mixed-species groups, and within-species populations, highlighting substantial genetic variation within and among species shaped by historical gene flow (particularly among diploid and sexual types), differences in ploidy levels, and adaptive divergence. The identification of genetically differentiated populations in wild species underscores their potential as reservoirs of unique adaptive traits such as drought tolerance, waterlogging and disease resistances, and nutrient-use efficiency. Meanwhile, the genetically diverse cultivated gene pool provides valuable resources for improving biomass productivity, nutritional quality, and resilience to environmental stress. Our findings provide a comprehensive genetic overview of one of the most important tropical forage genera and offer strategic insights for the conservation and utilization of *Urochloa* genetic resources.

## Introduction

Improving forage systems is essential for strengthening livestock feeds and global food and nutrition security. Enhanced forage systems also contribute to climate-resilient agriculture, support the restoration of degraded lands, and reduce greenhouse gas emission intensities (Bryan et al., 2013; Peters et al., 2013; Belay, 2019; Paul et al., 2020). Despite this importance, the livestock sector-particularly in sub-Saharan Africa (SSA)-continues to face persistent challenges in accessing adequate, high-quality feed resources (Mtengeti et al., 2008; Maleko et al., 2018). Among the tropical forages, *Urochloa* grass stands out as one of the most widely cultivated genera, with species well adapted to tropical and subtropical regions across the world (Mushagalusa et al., 2025; Worku et al., 2022; Ferreira et al., 2021).

The genus *Urochloa* (syn. *Brachiaria*) comprises approximately 100 species of tropical grasses widely distributed across the tropical and subtropical regions of the world, with several species playing a critical role as forages in livestock production systems (Higgins et al., 2022; Ferreira et al., 2021; Baptistella et al., 2020; Maass et al., 2015; Renvoize et al. 1996). Most species are polyploid, with ploidy levels ranging from diploid to hexaploid, and many reproduce apomictically, which is a form of asexual reproduction through seeds (Heslop-Harrison et al. 2023; Ferreira et al., 2021; van de Peer et al. 2021; Van De Peer et al. 2017). This poses both challenges and opportunities for genetic improvement. The genus exhibits considerable ecological and morphological diversity, with species adapted to a broad range of environments, from humid lowlands to drought-prone savannas (Jank et al., 2014; Pessoa-Filho et al., 2017). Among these, the cultivated species: *U. brizantha*, *U. decumbens*, *U. humidicola*, and *U. ruziziensis* underpin tropical pasture systems due to their high biomass yield, adaptability, and nutritive value, whereas wild species such as *U. mosambicensis*, *U. eminii*, and *U. oligotricha* harbour traits of potential agronomic importance, including drought tolerance, waterlogging resilience, and disease resistance (Higgins et al., 2022; Triviño et al., 2017).

The primary center of origin for *Urochloa* species is tropical Africa, where a rich diversity of wild germplasm exists (Vorontsova 2022; Ghimire et al. 2015; Morrone et al. 2012). From there, these grasses have been introduced and naturalized across tropical and subtropical regions of Latin America, Asia, and Oceania (Maass et al. 2015; Parsons 1972). Brazil has become a global leader in the cultivation and breeding of *Urochloa*, with millions of hectares planted to support its vast cattle industry. The Alliance of Bioversity-CIAT and EMBRAPA (Brazilian Agricultural Research Corporation) have developed the majority of cultivars that are now widely adopted worldwide (Ferreira et al., 2021; Jank et al. 2014). The wide ecological range of these species, spanning acidic soils, low-fertility lands, and drought-prone areas, makes them well-suited to marginal environments, thereby contributing significantly to sustainable forage systems.

The global collection of *Urochloa* species represents one of the most significant genetic resources for tropical forage improvement. The CGIAR conserves the largest collection of *Urochloa* species at the Alliance of Bioversity International and CIAT Genebank in Colombia and at the International Livestock Research Institute (ILRI) Forage Genebank in Ethiopia, primarily collected from SSA (Higgins et al., 2022; Ferreira et al., 2021). Additional collections are maintained by EMBRAPA in Brazil, Genetic Resources Research Institute (GeRRI) in Kenya, Australian Pastures Genebank (APG) in Australia, and the United States Department of Agriculture USDA in the United States (https://www.genesys-pgr.org/a/overview/v2QAYrExz0Q; Ferreira et al., 2021). These collections encompass key species including *U. brizantha, U. decumbens, U. humidicola,* and *U. ruziziensis*, which have been widely used in breeding programs for improved yield, stress tolerance, and environmental sustainability. Ongoing collaborative efforts focus on conserving wild relatives, characterizing genetic diversity, and identifying accessions suited to climate-resilient livestock systems in tropical and subtropical regions.

The high diversity and broad geographic distribution of *Urochloa* species present unique opportunities to investigate patterns of genetic differentiation, phylogeography, and adaptation. Recent advances in the generation of genome-wide markers and admixture-based clustering analyses enable the identification of distinct genetic lineages and subpopulations within and among species, providing insights into the processes of divergence, polyploidization, gene flow, and domestication (Masters et al., 2024; Higgins et al., 2022; Triviño et al., 2017). In particular, characterizing the population structure of wild species, such as *U. mosambicensis*, *U. jubata*, and *U. oligotricha,* can uncover novel alleles for traits of agronomic importance, including stress tolerance and nutritional quality, which are critical for improving forage productivity under climate variability (Masters et al., 2024; Higgins et al., 2022; Jank et al., 2014; Bastos et al., 2020).

In this study, we analysed 545 accessions representing 17 *Urochloa* species to investigate their genetic diversity, population structure, and phylogenetic relationships using cross-species genome-wide markers. We applied clustering and admixture analyses to define genetic lineages within the genus and examined the correspondence between genetic clusters and geographic origin, with the objectives of providing insights into the evolutionary relationships and potential genetic resources for breeding and conservation. The findings of this study offer a comprehensive framework for understanding the genetic architecture of *Urochloa* and provide a valuable foundation for the strategic utilization of germplasm in tropical forage improvement programs.

## Materials and methods

### Plant materials

This study included 545 accessions representing 17 *Urochloa* species, comprising both cultivated species (*U. brizantha, U. decumbens, U. humidicola,* and *U. ruziziensis*) and their wild relatives. These accessions were originally collected from 18 African countries and are maintained in the ILRI Forage Genebank at the Zway field site in Ethiopia. The commercial interspecific hybrid variety Mulato II, and a commercial cultivar of *U. decumbens,* Basilisk, were also included. Passport data, such as taxonomic classification, geographic origin, and accession codes, were compiled from the ILRI Genebank records (Supplementary File S1).

An additional 446 progeny plants, derived from seeds collected from 44 *U. brizantha* seed-parent accessions, were also included in the study. These progenies were evaluated to assess genetic relationships and segregation patterns among individuals originating from the same maternal parent and infer reproductive behaviour, and potential apomictic versus sexual reproduction within the accessions.

### DNA extraction and genotyping

Young leaf tissue from each plant was collected in a 2-ml Eppendorf tube, immediately placed in an icebox with ice, and transferred to a -80 °C freezer. Samples were freeze-dried for approximately 48 hours and ground into a fine powder using a tissue lyser. Genomic DNA was extracted from the powdered tissue using the DNeasy® Plant Mini Kit (Qiagen Inc., Valencia, CA) according to the manufacturer’s instructions. DNA concentration and quality were assessed using a NanoDrop spectrophotometer (DeNovix DS-11 FX) and 0.8% agarose gel electrophoresis. DNA concentrations were then adjusted to 50–100 ng/μl in 96-well semi-skirted plates and sent to Diversity Arrays Technology, Canberra, Australia for genotyping. Genotyping was performed using the DArTseq Genotyping-by-Sequencing (GBS) platform, which enables simultaneous marker discovery and genotyping of SNP and SilicoDArT markers by combining DArT complexity-reduction methods with next-generation sequencing (Muktar et al., 2019; Kilian et al., 2012).

### Mapping and filtering of markers

Markers were aligned with the 36 chromosomes of the *U. decumbens* genome (Ryan et al., 2025) using the short sequence fragments associated with each marker to determine their genomic positions. A heatmap showing the density and distribution of markers across the chromosomes was generated using an online tool available at https://www.bioinformatics.com.cn/en. For genetic diversity analysis, polymorphic markers were filtered based on the minor allele frequency (MAF), missing data thresholds, and pairwise linkage disequilibrium (LD) to ensure marker independence. These filtering steps were performed using the dartR (Gruber et al., 2018) and SNPRelate (Zheng et al., 2012) R-packages in R statistical software (R Core Team, 2022).

### Genetic integrity of plants within accessions

Genetic integrity of plants within an accession (within a plot) was assessed using Identity-by-descent (IBD) analysis based on the maximum likelihood (ML) relatedness estimator (Milligan 2003) using the R package SNPRelate (Zheng et al., 2012).

### Genetic diversity and population structure analysis

Genetic clustering was analysed using hierarchical clustering and principal component analysis (PCA) implemented in the Poppr (Kamvar et al., 2014) and adegenet (Jombart 2008) R-packages. Genetic diversity was estimated using pairwise Nei’s (Nei 1972) genetic distance. Population structure was conducted using hierarchical clustering and ADMIXTURE analysis (Alexander & Lange, 2011), which aimed to estimate individual ancestry proportions across a range of K values (from 2 to 20). Cross-entropy coefficients were plotted against K to determine the optimal number of ancestral populations, with the lowest cross-entropy value indicating the best-supported K (Frichot & François, 2015). Accessions with a membership coefficient (Q value) ≥ 0.60 for a particular cluster were classified as belonging predominantly to that population, while those with Q values < 0.60 were classified as admixtures (Gain & François, 2020). Genetic differentiation among clusters, sub-clusters, and collections was quantified using *F*_ST_, which ranges from 0 (no differentiation) to 1 (complete differentiation).

### Field phenotyping of *U. brizantha* accessions

Phenotypic data were collected from *U. brizantha* accessions maintained in conservation plots at the ILRI Forage Genebank, Zway field site, Ethiopia. At the onset of the rainy season, plants were uniformly cut to approximately 5 cm above ground level. Morphological traits, including plant height (PH, in cm), leaf length (LL, in cm), and leaf width (LW, in mm), were measured on three plants per accession and averaged. In addition, the presence or absence of leaf sheath hairs (LSH) and days to 50% flowering were recorded.

### Marker-trait associations

A genome-wide association study (GWAS) was performed using three statistical models implemented in the Genomic Association and Prediction Integrated Tool version 3 (GAPIT3) (Wang & Zhang, 2021): the General Linear Model (GLM), the Mixed Linear Model (MLM), and the SUPER model (Settlement of MLMs Under Progressively Exclusive Relationship). After filtering, a total of 106,053 markers (7,053 SNPs and 99,000 silicoDArTs) were retained. Marker filtering criteria included call rate (≥ 0.90), minor allele frequency (MAF ≥ 5%), missing data threshold (NA < 10%), and genome-wide distribution.

### Core collection

To identify a representative subset of accessions, we used the R package Core Hunter v3.2.1 (De Beukelaer et al., 2018). This tool selects core subsets by simultaneously optimizing multiple genetic criteria and allocation strategies, including overall genetic diversity, genetic clustering patterns derived from hierarchical clustering based on genome-wide SNP markers, and the geographical origins of the accessions. For assessing genetic diversity, we applied equal weighting to several metrics: modified Roger’s distance (RD), Shannon’s information index (SH), expected heterozygosity (EH), Shannon’s allelic diversity index (SH), and allele coverage (CV). These parameters were used collectively to define a core subset that represents the entire collection.

## Results

### GBS-based genotyping of *Urochloa* collections conserved at the ILRI Forage Genebank

We studied the genetic diversity of *Urochloa* spp. to enhance the utilization and efficient conservation of the species maintained in the ILRI Forage Genebank. From more than 580 *Urochloa* accessions representing 18 different species acquired since 1984 and maintained in the genebank, we successfully genotyped 566 accessions from 18 species using genotyping-by-sequencing (GBS) to assess genetic diversity within the *Urochloa* gene pool. The collection includes accessions obtained from 18 African countries (Supplementary file S1). Out of these initially genotyped 566 accessions, genotypes with high missing values for the genome wide markers were removed and a final set of 545 accessions from 17 species were studied. An additional 446 *U. brizantha* progeny plants were evaluated to assess genetic relationships and segregation patterns among individuals originating from the same seed parent, with the aim of inferring reproductive behaviour and distinguishing potential apomictic versus sexual reproduction within the accessions.

The GBS method of the DArTseq platform generated 150,478 SNP and 661,695 SilicoDArT genome-wide markers, of which 81% (121,871) and 35% (233,429) mapped onto the *U. decumbens* genome, respectively (Fig. 1). The percentages of the markers missing values ranged from 0 to 18% for the SilicoDArT markers and 0 to 80% for the SNP markers.

**Fig. 1.**
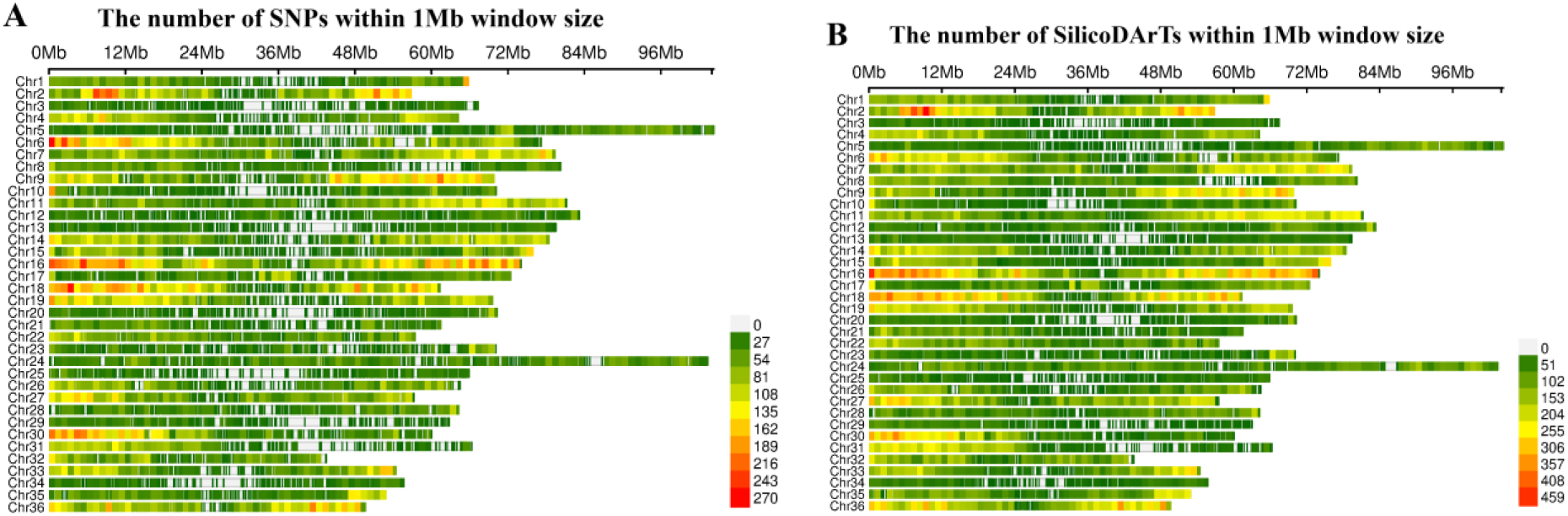
Heatmaps illustrating the genome-wide distribution and density of SNP (A) and SilicoDArT (B) markers across the 36 haploid chromosomes of the *U. decumbens* genome. Markers density was calculated within 1 Mb windows and is displayed by a colour gradient from green (low density), yellow (medium density), and red (high density).

The SNP markers heterozygosity (He) values ranged from 0 to 0.5 with an average value of 0.03 while polymorphic information content (PIC) values ranged from 0 to 0.5 with an average value of 0.14. For the SilicoDArT markers, for which PIC and He are equal, the values ranged from 0 to 0.5 with an average value of 0.07.

The markers missing values were substantially higher in the wild species compared to the species in the brizantha complex (*U. brizantha, U. decumbens*, and *U. ruziziensis*), which limited us to filter a sufficient number of genome-wide markers shared across species, particularly for SNP markers. Consequently, only 1% (2,044) of SNPs and 4% (24,271) of SilicoDArTs were filtered as common markers across the 17 species.

**Table 1.**
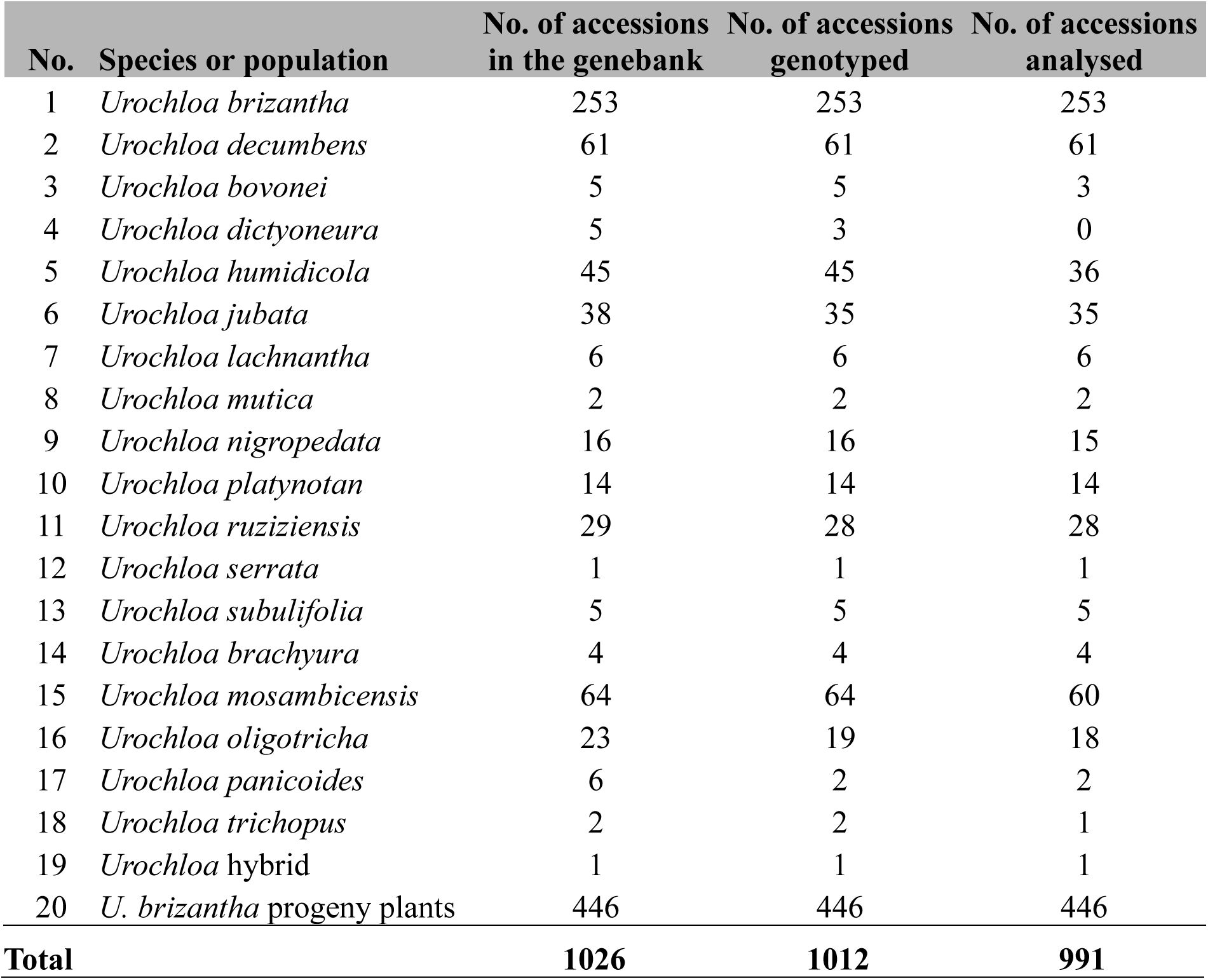
*Urochloa* species included in this study, with the number of accessions per species maintained in the ILRI Forage Genebank and the number of accessions genotyped and analysed. Detailed information for each accession is provided in Supplementary file S1.

### Genetic diversity and clustering across *Urochloa* species

Before genetic diversity analysis, we first examined the genetic integrity of plants within accessions, by genotyping two to three plants per accession, based on a pairwise IBD (Identity-By-Descent) analysis using the Maximum Likelihood Estimation (MLE). Based on the IBD analysis, we retained 545 accessions that were considered true-to-type (Supplementary file S2). Where multiple true-to-type individuals were found in an accession, the individual with the least missing data was selected for diversity analysis.

A filtered marker set of 2,044 SNPs and 24, 271 SilicoDArTs (15% missing values, minor allele frequency of 0.3%, and a maximum linkage-disequilibrium value of 0.7) were used for clustering analysis. Hierarchical clustering based on Nei’s genetic distance grouped the 545 accessions of 17 *Urochloa* species into two major groups (Groups I and II) with a 100% bootstrap value, which revealed a clear division between the brizantha complex (*U. brizantha, U. decumbens,* and *U. ruziziensis,* including Mulato-II-the interspecific hybrid commercial variety) and the remaining fourteen species (*U. mosambicensis, U. panicoides, U. trichopus, U. brachyura, U. platynota, U. oligotricha, U. serrata, U. nigropedata, U. mutica, U. lachnantha, U. humidicola, U. jubata, U. subulifolia,* and *U. bovonei*), which are primarily wild and less domesticated species (Fig. 2a).

**Fig. 2.**
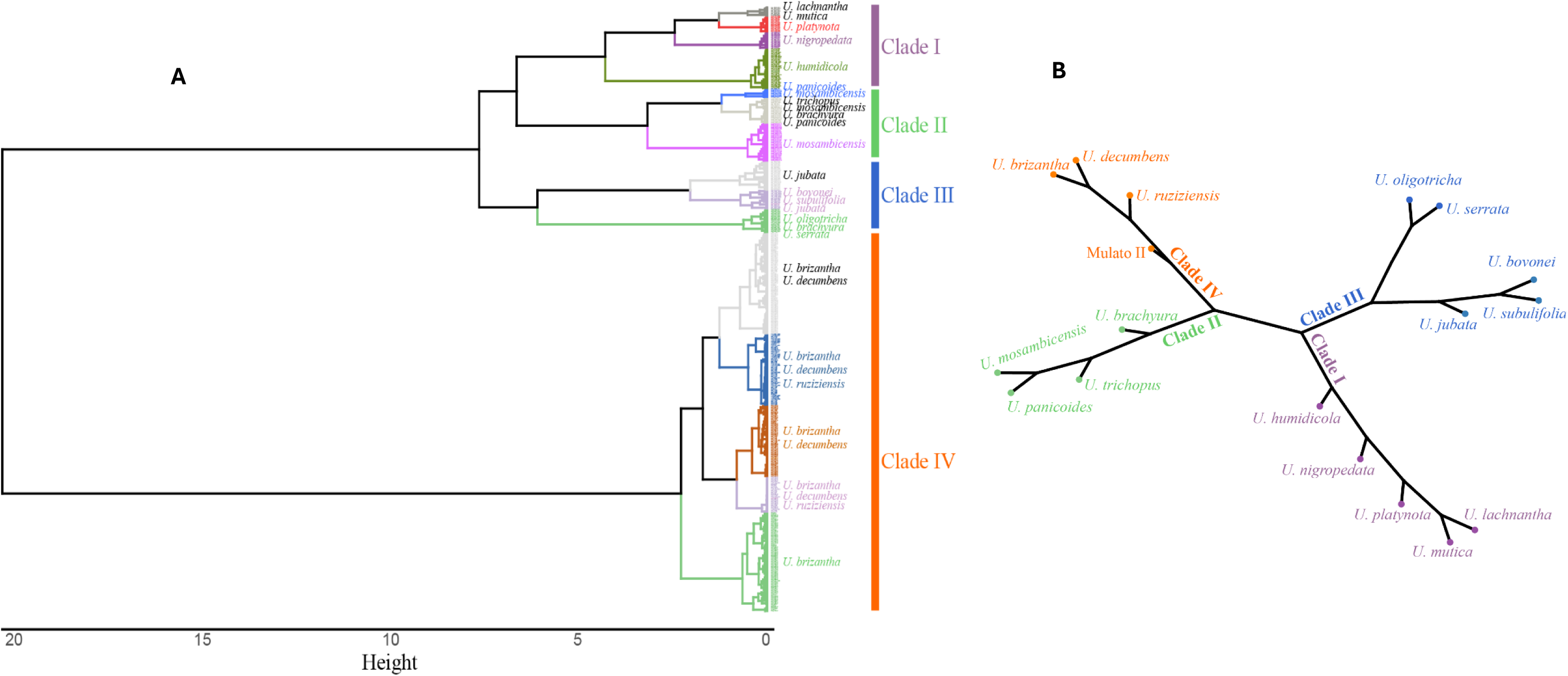
Phylogenetic trees based on hierarchical clustering showing groups and clusters of the entire collection of *Urochloa* accessions and the corresponding species analysed in the study. (**A**) Hierarchical clustering for all individual accessions, suggesting two major groups and 15 subclusters. (**B**) Hierarchical clustering for species, after categorizing the accessions according to their respective species, suggesting the four major clades.

The mean Nei’s D values for Groups I and II were 0.13 and 0.31, respectively, indicating greater genetic diversity in group II, which includes most of the wild species. This higher diversity was also evident from the longer branch lengths observed in the dendrogram. The two major groups, Groups I and II, further sub-grouped into five and ten clusters, respectively (Fig. 2a).

The genetic relationships among the 17 species were analysed using hierarchical clustering based on Nei’s genetic distance, after categorizing the accessions according to their respective species. The analysis revealed four distinct clusters, likely representing different clades. Clade-I comprised *U. humidicola, U. lachnantha, U. Mutica, U. Platynota,* and *U. Nigropedata*. Clade-II included *U. mosambicensis, U. panicoides, U. Trichopus,* and *U. brachyura*. Clade-III contained *U. jubata, U. subulifolia, U. bovonei, U. oligotricha,* and *U. serrata*. Clade-IV grouped the brizantha complex - *U. brizantha, U. decumbens,* and *U. ruziziensis,* together with Mulato-II-a commercial interspecific hybrid variety (Fig. 2b). Mean genetic distance (Nei’s D) values ranged from 0.13 to 0.47, with an overall average of 0.27. Clade III exhibited the highest mean Nei’s D (0.47), followed by Clade I (0.37) and Clade II (0.15), while Clade IV showed the lowest value (0.13).

### Admixture analysis

We conducted an admixture analysis to define clusters (populations) within each of the four clades, estimating between three to five clusters based on CV error values (Supplementary Figure S2). Applying a minimum membership probability threshold of 60%, the 73 accessions in Clade-I were assigned to five distinct clusters, with no admixture group, each precisely corresponding to one of the five species within the clade (Fig. 3). Specifically, all 36 *U. humidicola* accessions were assigned to cluster-I, all 14 *U. platynotan* accessions to Cluster-II, six *U. lachnantha* accessions to cluster-III, 15 *U. nigropedata* accessions to Cluster-IV, and two *U. mutica* accessions to Cluster-V. Pairwise *F*_ST_ among the five clusters varied from 0.75 between Clusters I and V (between *U. humidicola* and *U. mutica*) to 0.94 between Clusters II and III (between *U. platynotan* and *U. lachnantha*). With the exception of Cluster-I, the clustering pattern appears to reflect the geographic origin of the accessions (Fig. 4). The accessions in Cluster-II, which contains *U. platynota*, were from Rwanda and Burundi. All *U. lachnantha* accessions in Cluster-III originated from Ethiopia, while all *U. nigropedata* accessions in Cluster-IV were from Zimbabwe. The *U. humidicola* accessions in Cluster-I were distributed across East African countries, including Ethiopia, Tanzania, Burundi, and Zambia (Fig. 4). In contrast, the geographic origin of the two *U. mutica* accessions was unknown.

**Fig. 3.**
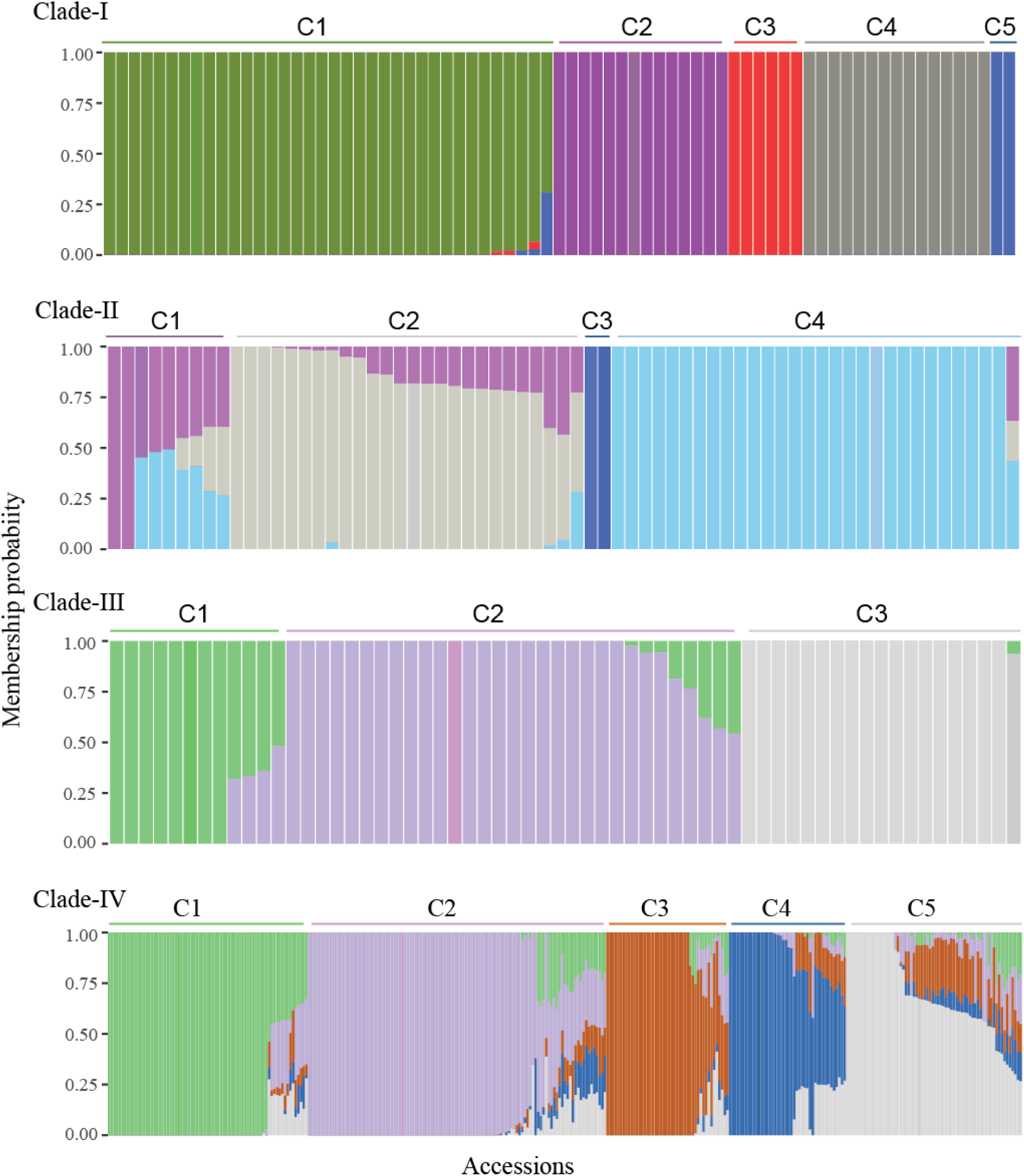
Bar plots of admixture analysis showing the inferred genetic ancestry within each of the four clades (I–IV). Each accession is shown as a vertical bar partitioned according to the proportional contribution of ancestral genetic clusters. Accessions represented by a single colour indicate ‘pure’ ancestry, whereas those with two or more colours reflect admixture from multiple ancestral sources. Each ancestral cluster is depicted by a distinct colour. A minimum ancestral membership probability threshold of 60% was applied to assign accessions to a cluster.

**Fig. 4.**
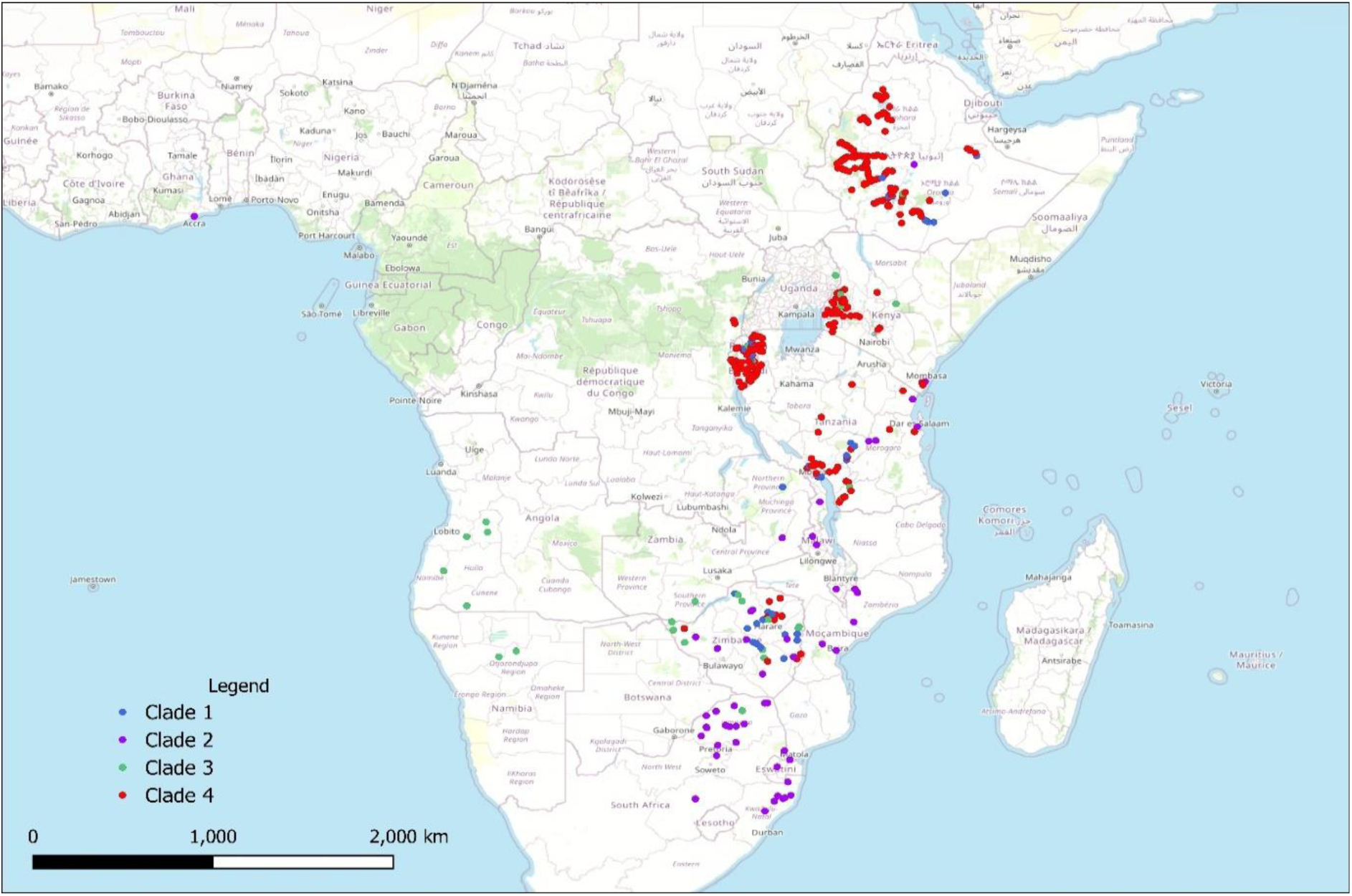
A map illustrating the spatial distribution and geographical origin of the 545 *Urochloa* accessions used in the study across Africa. Each point represents an accession, colour-coded according to its assigned clade: blue for Clade I, purple for Clade II, green for Clade III, and red for Clade IV.

The 67 accessions in Clade-II, representing four species (*U. mosambicensis*, *U. panicoides*, *U. trichopus*, and *U. brachyura*), were divided into four clusters and an admixture group (Fig. 3). Clusters-I and II comprised two and 23 *U. mosambicensis* accessions, respectively; Cluster-III contained two *U. brachyura* accessions, while Cluster-IV consisted of 29 accessions from *U. mosambicensis* (24), *U. brachyura* (2), *U. panicoides* (2), and *U. trichopus* (1). Pairwise *F_ST_* among the four clusters varied from 0.24 between Clusters-II and IV to 0.88 between Clusters-I and III as well as between Clusters-III and IV. Accessions within species were distributed across different clusters, representing distinct populations. Accordingly, the 60 *U. mosambicensis* accessions divided into three populations: Population-I (2 accessions), Population-II (23 accessions), and Population-III (24 accessions), with 11 accessions remaining admixed. Similarly, two populations, with two accessions in each, were detected in *U. brachyura*. While most accessions of the four species in Clade-II originated from Southern and Southeastern Africa, the majority of *U. mosambicensis* accessions in Population-I were from South Africa, whereas those in Population-II were from Southeastern African countries, including Mozambique, Zimbabwe, Malawi, and Tanzania (Fig. 4).

The admixture analysis classified the 62 accessions in Clade-III into three distinct clusters, along with an admixture group. Cluster-I comprised a mix of three species: *U. subulifolia* (5 accessions), *U. bovonei* (4 accessions), and *U. jubata* (2 accessions). Cluster-II consisted of 29 accessions purely from *U. jubata*, with 3 accessions exhibiting shared ancestry with Cluster-I. All *U. oligotricha* (18 accessions) were in Cluster-III, showing no shared ancestry with other clusters. Additionally, the single accession of *U. serrata* was grouped within this cluster. Pairwise *F_ST_* values among the three clusters ranged from 0.35 between Clusters-I and II to 0.90 between Clusters-II and III. The 35 *U. jubata* accessions were grouped into two populations, with the majority assigned to Population-II. Population I consisted of three accessions that showed shared ancestry with Population-II, while an additional three accessions were classified as an admixed group. Cluster-II predominantly comprised accessions originating from East African countries, including Ethiopia, Kenya, Rwanda, and Burundi. In contrast, the majority of accessions in Clusters-I and III were derived from Southern African countries, such as Zimbabwe and Angola (Fig. 4).

Clade IV, which comprises the highest number of accessions from the three known species in the brizantha complex, was subdivided into three major clusters and five subclusters along with an additional admixture group. Subcluster-I predominantly consisted of *U. ruziziensis* (24 accessions, accounting for 86%) and *U. decumbens* (30 accessions, or 48%), along with a small representation from *U. brizantha* (6 accessions, or 2%). This subcluster exhibited no evidence of shared ancestry with the other subclusters, indicating a genetically distinct group. Subclusters-II and IV were composed exclusively of *U. brizantha* accessions. Subcluster-II represented 86 accessions, or approximately 34% of the total, while Subcluster-IV contained 27 accessions, making up around 11%. Subcluster-III was a more heterogeneous group, comprising accessions from all three species. It included 12 accessions of *U. decumbens* (20%), 4 accessions of *U. ruziziensis* (14%), and 19 accessions of *U. brizantha* (8%), reflecting a mixed ancestry. Subcluster-V also represented a combination of species, primarily *U. brizantha* (39 accessions, or 15%) and *U. decumbens* (7 accessions, or 11%). The admixture group contained 89 accessions in total, made up of 76 accessions of *U. brizantha* (30%), 12 accessions of *U. decumbens* (20%), and the single accession of Mulato II, a commercial interspecific hybrid variety. This group suggests a history of gene flow among species and populations and possibly targeted breeding efforts. Pairwise *F_ST_* values among the five subclusters ranged from 0.15 between Subclusters-III and V to 0.84 between Subclusters-I and II, indicating substantial genetic differentiation, particularly between subclusters composed of different species. Accessions from each of the three species were distributed across multiple clusters or subclusters, representing genetically distinct populations. Accordingly, three major and five sub populations were identified within *U. brizantha*, three within *U. decumbens*, and two within *U. ruziziensis*. The majority of accessions from the three species within Clade-IV originated from Eastern African countries, particularly Ethiopia, Kenya, Burundi, and Rwanda (Fig. 4). The majority of *U. brizantha* accessions originated from Ethiopia, followed by Kenya and Burundi. In contrast, *U. decumbens* accessions were primarily collected from Rwanda and Kenya, with additional contributions from Burundi. For *U. ruziziensis*, most accessions came from Burundi, followed by Rwanda and Kenya.

### Genetic diversity in *U. brizantha* progeny plants

Of the 545 accessions representing 17 species included in this study, over 46% belonged to *U. brizantha*. Consequently, we conducted further analyses focusing specifically on this species. Genetic relationship analysis among progeny plants derived from the same seed parent, using identity by descent (IBD) and Nei’s genetic distance, revealed no detectable genetic variation among progeny originating from the same seed parent across the 44 *U. brizantha* accessions. This uniformity suggests that the seeds were likely produced either through apomixis, an asexual reproductive process that results in clonal offspring, or self-pollination, which limits genetic recombination. While neither of these reproductive mechanisms introduces new genetic variation, they are essential for maintaining the genetic integrity and uniformity of the accessions-ensuring that they remain true-to-type across generations. Principal Component Analysis (PCA) identified three distinct genetic clusters. Interestingly, seed parents were distributed among these clusters together with their corresponding progeny (Fig. 5), indicating strong genetic similarity and confirming the lack of genetic variation between the progeny and the same seed parent observed by the IBD analysis. This further supports the conclusion that the progenies were derived through apomictic or self-pollinated reproduction, resulting in minimal genetic divergence from the seed parent.

**Fig. 5.**
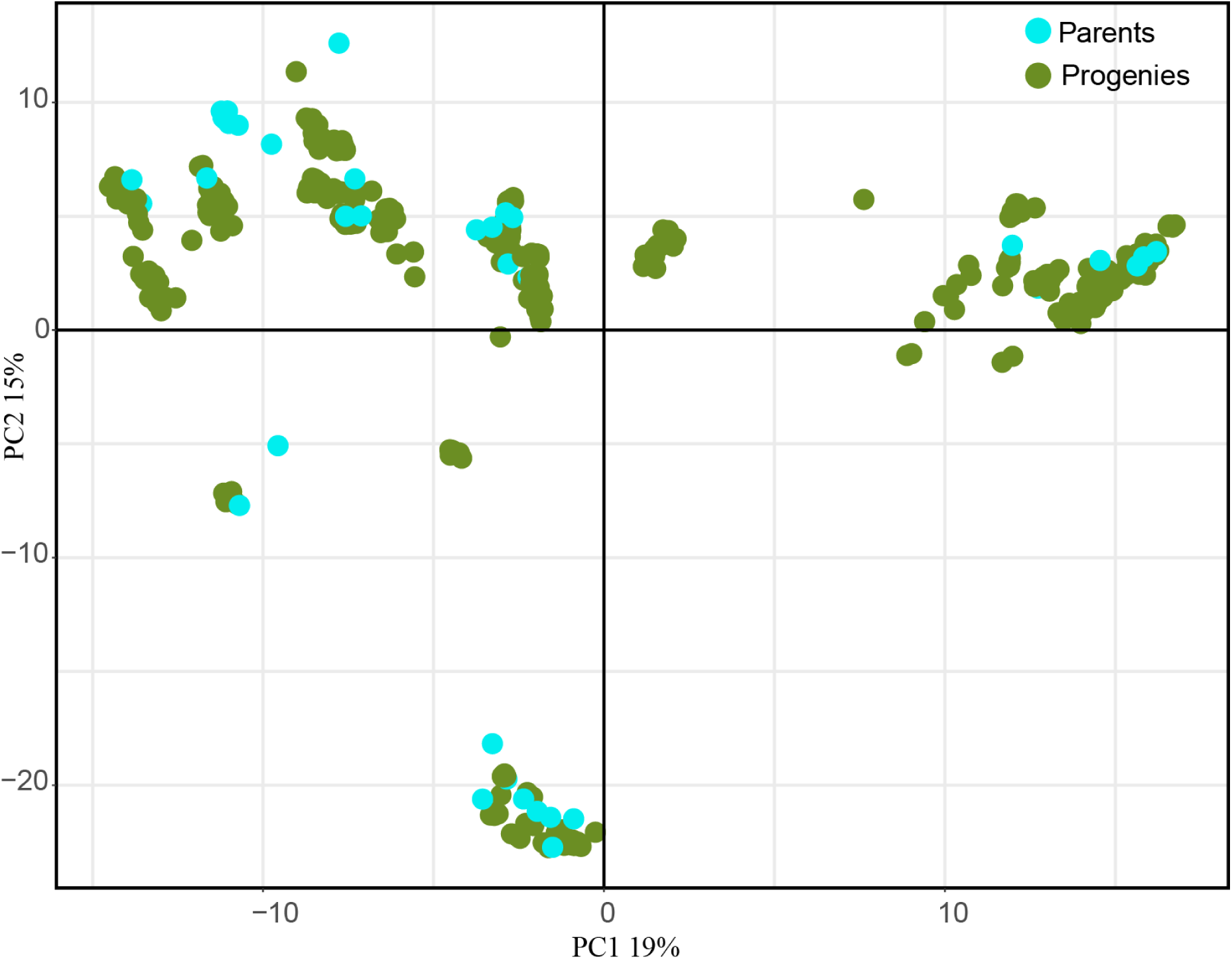
Principal component analysis (PCA) showing population structure and the distribution of 446 progeny plants (coloured blue) derived from 44 *U. brizantha* seed parents (coloured green).

### Phenotypic variation among genetic clusters and marker-trait associations in *U. brizantha* collection

Phenotypic data for eight traits collected from 241 *U. brizantha* accessions maintained in the conservation plots at the ILRI Zway site were used to assess correlations between trait variation and the genetic clusters identified by the genetic diversity analysis. The results revealed notable associations, particularly for days to flowering, leaf length, plant height, and number of spikelet rows (Fig. 6A). While each cluster exhibited some degree of phenotypic variation, Cluster-2 was characterized by late-maturing, taller accessions with longer leaves. In contrast, Clusters-1 and 5 consisted mainly of early-maturing accessions with shorter height and smaller leaves.

**Fig. 6.**
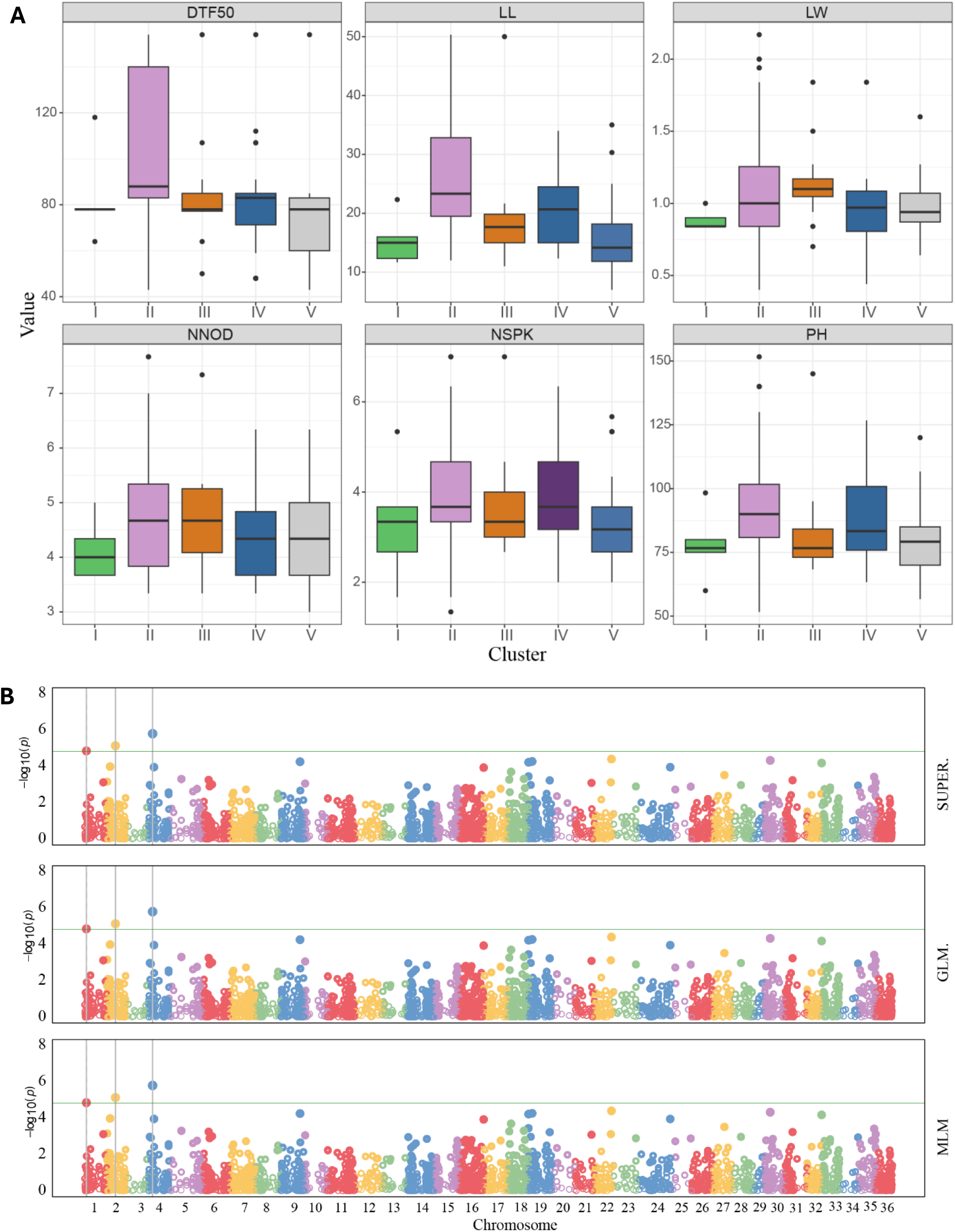
Molecular characterization of the *U. brizantha* accessions**. (A)** Boxplots showing correlation between trait variations and genetic clusters. The colours are according to the ADMIXTURE analysis in Clade IV. **(B)** Manhattan plot showing the distribution of significant marker-trait associations identified for leaf sheath hair. The vertical axis at the left represents the -log10 *p*-value, and markers and their chromosome positions are shown on the horizontal axis. The three GAPIT models (MLM, GLM, SUPER) used are shown on the right vertical axis.

Significant associations were identified for leaf sheath hair, leaf length, and days to 50% flowering (Fig. 6B). Notably, the association with leaf sheath hair was consistently detected in both SNP and SilicoDArT markers, in separate analyses, explaining 7–24% of the phenotypic variation. These markers represent valuable resources for the genetic improvement of *U. brizantha* through marker-assisted selection and will also support further characterization and map-based cloning of the underlying QTLs.

### A core collection for the *U. brizantha* collection

To facilitate efficient conservation, distribution and utilization of the *U. brizantha* genetic resource with 253 accessions, a core collection comprising 51 accessions was established. This core set was carefully selected to capture the maximum possible diversity present within the entire *U. brizantha* collection. The selection process integrated multiple sources of information, including a subset of informative genome-wide SNP markers, the geographical origins of the accessions, and the clustering patterns derived from hierarchical clustering analysis. By combining these criteria, the resulting core collection represents a genetically and geographically diverse subset that can serve as a valuable resource for future research, breeding, and conservation efforts.

The core collection exhibited greater genetic diversity compared to the entire germplasm collection, as evidenced by multiple diversity indices. When assessed using the modified Rogers (MR) distance, the average genetic distance within the core was 0.29, nearly double that of the whole collection (MR = 0.15), indicating a broader spectrum of genetic variation among the selected accessions. Similarly, other key diversity metrics, including expected heterozygosity (EH) and Shannon’s allelic diversity index (SH), were consistently higher in the core, further supporting its enriched genetic variability. The core collection fully captured the genetic structure of the whole collection. All population clusters identified in the whole collection were retained in the core, confirming its representativeness (Fig. 7; Table 2).

**Fig. 7.**
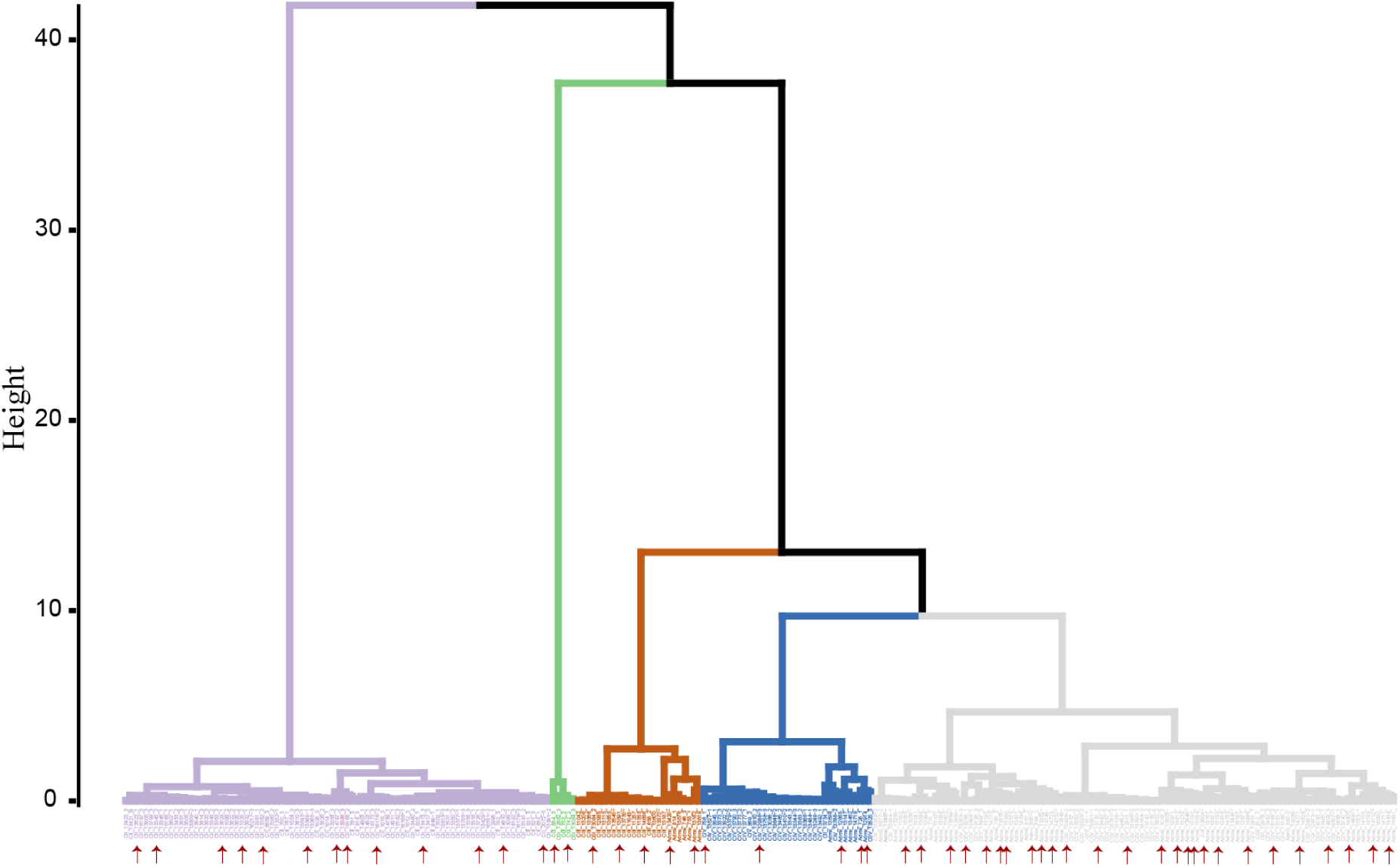
Hierarchical clustering depicting the three major and five sub populations detected in the entire 253 *U. brizantha* accessions. The positions of the 51 core subset accessions are shown by arrows.

**Table 2.** The 51 accessions selected as core subset representing the genetic diversity of the entire *U. brizantha* collection maintained in the ILRI Forage Gnebank.

| No. | Acc. | DOI | Species | Origin | Ploidy | Cluster |
| --- | --- | --- | --- | --- | --- | --- |
| 1 | 1019 | 10.18730/FP8VZ | *Urochloa brizantha* | Burundi | NA | Admx |
| 2 | 12865 | 10.18730/FRQK= | *Urochloa brizantha* | Kenya | NA | Admx |
| 3 | 12914 | 10.18730/FRS4A | *Urochloa brizantha* | Kenya | NA | Admx |
| 4 | 13055 | 10.18730/FRVMG | *Urochloa brizantha* | Kenya | NA | Admx |
| 5 | 13099 | 10.18730/FRWPD | *Urochloa brizantha* | Kenya | NA | Admx |
| 6 | 13182 | 10.18730/FRYN2 | *Urochloa brizantha* | Kenya | NA | Admx |
| 7 | 13192 | 10.18730/FRYW9 | *Urochloa brizantha* | Kenya | 4 | Admx |
| 8 | 13434 | 10.18730/FS4WG | *Urochloa brizantha* | Ethiopia | NA | Admx |
| 9 | 13762 | 10.18730/FSECR | *Urochloa brizantha* | Ethiopia | NA | Admx |
| 10 | 14698 | 10.18730/FT36R | *Urochloa brizantha* | Rwanda | NA | Admx |
| 11 | 14725 | 10.18730/FT3W9 | *Urochloa brizantha* | Rwanda | 5 | Admx |
| 12 | 13130 | 10.18730/FRXDU | *Urochloa brizantha* | Kenya |  | V |
| 13 | 14777 | 10.18730/FT59H | *Urochloa brizantha* | Burundi | 4 | Admx |
| 14 | 14800 | 10.18730/FT5WU | *Urochloa brizantha* | Burundi | 4 | Admx |
| 15 | 14812 | 10.18730/FT68B | *Urochloa brizantha* | Burundi | NA | Admx |
| 16 | 726 | NA | *Urochloa brizantha* | NA | NA | Admx |
| 17 | 810 | NA | *Urochloa brizantha* | NA | NA | Admx |
| 18 | 14712 | 10.18730/FT3H= | *Urochloa brizantha* | Rwanda | NA | Admx |
| 19 | 12913 | 10.18730/FRS39 | *Urochloa brizantha* | Kenya |  | V |
| 20 | 13352 | 10.18730/FS2TR | *Urochloa brizantha* | Ethiopia | 4 | Admx |
| 21 | 13594 | 10.18730/FS9FF | *Urochloa brizantha* | Ethiopia | 4 | Admx |
| 22 | 13646 | 10.18730/FSAZT | *Urochloa brizantha* | Ethiopia | 5 | Admx |
| 23 | 880 | NA | *Urochloa brizantha* | NA | NA | Admx |
| 24 | 13420 | 10.18730/FS4K7 | *Urochloa brizantha* | Ethiopia | NA | Admx |
| 25 | 13370 | 10.18730/FS3B4 | *Urochloa brizantha* | Ethiopia | 4 | Admx |
| 26 | 13380 | 10.18730/FS3NE | *Urochloa brizantha* | Ethiopia | 4 | Admx |
| 27 | 742 | NA | *Urochloa brizantha* | NA | NA | I |
| 28 | 13256 | 10.18730/FS0DN | *Urochloa brizantha* | Kenya | NA | V |
| 29 | 14809 | 10.18730/FT658 | *Urochloa brizantha* | Burundi | NA | V |
| 30 | 16189 | 10.18730/FVCNK | *Urochloa brizantha* | Zimbabwe | NA | V |
| 31 | 13734 | 10.18730/FSDKU | *Urochloa brizantha* | Ethiopia | 4 | II |
| 32 | 14787 | 10.18730/FT5GR | *Urochloa brizantha* | Burundi | NA | II |
| 33 | 16168 | 10.18730/FVC31 | *Urochloa brizantha* | Zimbabwe | NA | II |
| 34 | 13591 | 10.18730/FS9DD | *Urochloa brizantha* | Ethiopia |  | III |
| 35 | 14686 | 10.18730/FT2XF | *Urochloa brizantha* | Burundi | NA | III |
| 36 | 13604 | 10.18730/FS9SS | *Urochloa brizantha* | Ethiopia | 4 | III |
| 37 | 12986 | 10.18730/FRV1' | *Urochloa brizantha* | Kenya | 4 | IV |
| 38 | 13345 | 10.18730/FS2NK | *Urochloa brizantha* | Ethiopia | NA | II |
| 39 | 13629 | 10.18730/FSAHC | *Urochloa brizantha* | Ethiopia | 4 | II |
| 40 | 13382 | 10.18730/FS3QG | *Urochloa brizantha* | Ethiopia | 4 | II |
| 41 | 13369 | 10.18730/FS3A3 | *Urochloa brizantha* | Ethiopia | 4 | II |
| 42 | 16176 | 10.18730/FVCB9 | *Urochloa brizantha* | Zimbabwe | NA | II |
| 43 | 13413 | 10.18730/FS4D1 | *Urochloa brizantha* | Ethiopia | 4 | II |
| 44 | 2072 | 10.18730/FZH87 | *Urochloa brizantha* | Ethiopia | NA | II |
| 45 | 884 | NA | *Urochloa brizantha* | NA | NA | I |
| 46 | 13084 | 10.18730/FRW90 | *Urochloa brizantha* | Kenya | NA | I |
| 47 | 756 | NA | *Urochloa brizantha* | NA | NA | IV |
| 48 | 13436 | 10.18730/FS4YJ | *Urochloa brizantha* | Ethiopia | 5 | V |
| 49 | 13623 | 10.18730/FSAB6 | *Urochloa brizantha* | Ethiopia | 4 | II |
| 50 | 11335 | 10.18730/FQ7NJ | *Urochloa brizantha* | Ethiopia | 4 | II |
| 51 | 13753 | 10.18730/FSE4G | *Urochloa brizantha* | Ethiopia | NA | Admx |

## Discussion

In this study, we analysed the genetic diversity existing in the extensive *in*-*situ* collection of *Urochloa* spp. maintained in the ILRI Forage Genebank for the last four decades at its Zway field site in Ethiopia. The collection contains accessions collected from 18 different countries, the majority originating from Africa, and represent 17 distinct species, including several key pasture species of global importance, such as *U. brizantha, U. decumbens, U. ruziziensis, U. humidicola,* and *U. mosambicensis*.

We genotyped the collection using the GBS method of the DArTseq platform that generated a large number of SNP and SilicoDArT genome-wide markers, providing valuable resources for genetic analysis in *Urochloa* spp. A substantial proportion of missing data was observed, particularly among SNPs in the wild *U.* species. This pattern is consistent with previous reports that wild relatives often yield lower-quality SNP datasets due to higher genetic divergence from the cultivated species (Mushagalusa et al., 2025; Ferreira et al., 2021), which mostly used in selection of representative fragments of the gene pool in GBS based sequencing technology (https://www.diversityarrays.com). The high rate of missing SNPs in the wild accessions consequently reduced the number of markers shared across species, thereby limiting the scope of cross-species genetic diversity analyses. These findings highlight both the potential and the challenges of applying genotyping-by-sequencing platforms in highly diverse germplasm collections.

Unlike the SNPs, SilicoDArTs (presence/absence variants) showed higher retention rates and provided a broader and more uniform representation of the *Urochloa* genome across both cultivated and wild species, making them more robust for genotyping genetically diverse germplasm as compared to SNPs from the platform. Nevertheless, we were able to filter a set of genome-wide SNP and SilicoDArT markers suitable for cross-species analysis across the *Urochloa* collection.

### Phylogenetic clades and evolutionary relationships within the *Urochloa* genus

Hierarchical clustering analysis using the filtered markers provided clear resolution among the 545 accessions representing 17 *Urochloa* species, revealing distinct patterns of genetic differentiation. The resulting clustering separated the accessions into two major groups that clearly distinguished the three significant species of the brizantha complex (*U. brizantha, U. decumbens,* and *U. ruziziensis*) from the rest of the species mainly composed of wild and less domesticated relatives. This separation highlights the strong genetic divergence between domesticated and wild gene pools, likely reflecting the effects of selection during domestication and adaptation (particularly in the key forage species *U. brizantha* and *U. decumbens*), shared ancestry, and historical polyploidization events (Higgins et al., 2022; Ferreira et al., 2021; Pessoa-Filho et al., 2017). Recent studies indicate that *U. ruziziensis* is likely the ancestral donor of one of the sub-genomes (the B² sub-genome) in tetraploid/allopolyploid *U. decumbens* and may also have contributed to the complex evolutionary origin of *U. brizantha* (Ryan et al., 2025; Ferreira et al., 2021; Pessoa-Filho et al., 2017). Furthermore, the analysis identified four major clusters corresponding to four well-defined phylogenetic clades, mostly consistent with previously reported evolutionary relationships within the genus (Masters et al., 2024; Renvoize et al., 1996; Triviño et al., 2017).

Clade I comprised *U. humidicola, U. lachnantha, U. mutica, U. platynota*, and *U. nigropedata*. The clustering of *U. humidicola* and *U. platynota* together was consistent with a previous report (Renvoize et al., 1996), suggesting a possible shared ancestry associated with adaptation to waterlogged or moist environments. However, this clustering does not correlate with the report of Masters et al. (2024), in which *U. mutica* and *U. platynotan* were placed in distinct clades. Members of this clade are generally recognized for their tolerance to waterlogging and ability to thrive under high soil moisture conditions, traits that make them particularly valuable for pasture systems in humid regions (Villegas et al., 2022; Ferreira et al., 2021; Renvoize et al., 1996). An admixture analysis further resolved this clade into five clearly differentiated genetic clusters, each aligning exactly with one of the five species in the clade and showing no evidence of significant admixture. The very high *F*_ST_ values among clusters, together with the strong correspondence between genetic clusters and species identity, indicate limited introgression among these taxa, possibly due to differences in ploidy level, reproductive mode, or ecological niches (Higgins et al., 2022; Triviño et al., 2017). The observed geographic structuring further highlights the role of regional isolation in shaping genetic divergence among the species within the clade. For example, the concentration of *U. lachnantha* in the Ethiopian highlands and *U. nigropedata* in oSuthern Africa may have led to divergent selection pressures, driving local adaptation and reproductive isolation. In contrast, the broad distribution of *U. humidicola* across multiple countries may explain its greater intra-specific diversity and widespread ecological success in tropical lowlands (Tomaszewska et al., 2023; Villegas et al., 2022). Collectively, the admixture results provide valuable insights into the evolutionary and ecological processes that have shaped Clade I, indicating that both geography and reproductive mechanisms have played central roles in maintaining genetic distinctiveness among its constituent species. Furthermore, the genetic distinctiveness of Clade I from the cultivated brizantha complex highlights its potential as a source of novel alleles for improving stress tolerance and biomass productivity in breeding programs targeting marginal or waterlogged environments (Villegas et al., 2022; Renvoize et al., 1996).

Clade II comprised of *U. mosambicensis, U. panicoides, U. trichopus,* and *U. brachyura*. Their clustering aligns with previous reports (Masters et al., 2024; dos Santos Pessoa et al. 2022; Renvoize et al., 1996), which describe these species as typically distributed across the arid and semi-arid regions of sub-Saharan Africa adapting to hot and drought-prone environments. Their genetic distinctiveness from the humid-adapted species of Clade I likely reflects divergent evolutionary routes shaped by ecological adaptation and geographic isolation. The genetic variation within this clade thus represents a valuable reservoir of alleles that could be exploited to enhance the resilience and productivity of improved forage cultivars in water-limited environments. The admixture analysis revealed substantial population structure within this clade, dividing it into four distinct genetic clusters and one admixture group. Clusters I and II were composed exclusively of *U. mosambicensis* accessions, while Cluster III contains *U. brachyura* accessions. Cluster IV represented a mixed group from the four species: *U. mosambicensis*, *U. brachyura*, *U. panicoides*, and *U. trichopus*, indicating a degree of shared ancestry or historical gene flow among these taxa. The occurrence of admixed individuals and the clustering of multiple species in Cluster IV suggest possible hybridization events among *U. mosambicensis, U. brachyura,* and *U. panicoides*, potentially facilitated by overlapping flowering times and geographic distributions (Renvoize et al., 1996). The presence of genetically admixed accessions further highlights the dynamic evolutionary history of this clade, shaped by both divergence and occasional introgression.

Clade III composed of five species: *U. jubata, U. subulifolia, U. bovonei, U. oligotricha,* and *U. serrata*, a grouping mostly consistent with previous phylogenetic studies (Masters et al., 2024, Triviño et al., 2017). The species are primarily distributed across East and Southern Africa and represent a more diverse group of wild species with limited domestication history. Their genetic uniqueness highlights their potential as untapped resources for introducing novel traits such as pest resistance and adaptation to marginal soils (Tomaszewska et al., 2023; Valério et al., 1996). Admixture analysis further revealed substantial genetic differentiation within Clade III, partitioning the accessions into three well-defined clusters alongside an admixture group. Cluster I consisted of a mixed species composition, including *U. subulifolia, U. bovonei,* and *U. jubata*, suggesting possible shared ancestry. Although historical hybridization among these taxa cannot be ruled out, it is likely limited due to differences in ploidy level, reproductive mode, and ecological separation. In contrast, Cluster II and Cluster III contained accessions exclusively from *U. jubata* and *U. oligotricha*, respectively. Cluster II was mainly composed of accessions from East African countries, including Ethiopia, Kenya, Rwanda, and Burundi. In contrast, the majority of accessions in Clusters I and III originated from Southern Africa, particularly Zimbabwe and Angola. This north–south differentiation likely reflects environmental gradients and historical dispersal barriers, consistent with the biogeographic differentiation observed across other *Urochloa* taxa (Masters et al., 2024; Higgins et al., 2022). From a practical standpoint, the marked genetic differentiation observed among Clade III species and populations underscores their potential as reservoirs of allelic variation for forage improvement. In particular, *U. jubata* and *U. oligotricha* may provide valuable genetic resources for enhancing traits such as tolerance to drought and low soil fertility, important factors for sustainable livestock feed production in tropical and subtropical regions.

Clade IV comprised of the largest number of accessions, representing the three well-known species of the brizantha complex: *U. brizantha, U. decumbens,* and *U. ruziziensis*. These species (including *U. humidicola*) form the core of tropical pasture systems across Africa, Latin America, and Asia due to their high biomass yield, adaptability, and nutritional value (Mushagalusa et al., 2025; Higgins et al., 2022; Ferreira et al., 2021; Triviño et al., 2017). The close genetic relatedness among these species reflects their shared evolutionary ancestry, historical polyploidization events, and selection pressures associated with domestication (Ryan et al., 2025; Ferreira et al., 2021; Pessoa-Filho et al., 2017). They have been extensively used in breeding programs aimed at combining the high productivity of *U. brizantha*, the adaptation to acidic and low fertile soil of *U. decumbens*, and the sexual fertility, high seed production, and nutritional quality of *U. ruziziensis* (Ferreira et al., 2021; Jank et al., 2014). These findings reinforce previous reports that the brizantha complex forms a distinct and relatively narrow genetic group within *Urochloa*, resulting from both natural and artificial selection (Triviño et al., 2017; Ferreira et al., 2016), and highlights the need to broaden its genetic base through targeted introgression from other clades. The admixture analysis revealed a complex population structure within Clade IV, identifying three major clusters, five subclusters, and an additional admixture group. Subcluster I was predominantly composed of *U. ruziziensis* (86%) and *U. decumbens* (48%), with minor representation from *U. brizantha* (2%). This minor grouping of *U. brizantha* accessions within this subcluster could be attributed to either initial errors in taxonomic classification or to contamination that occurred over the course of long-term conservation in the genebank plots. This subcluster exhibited no evidence of shared ancestry with other subclusters, indicating that it represents a genetically distinct lineage within Clade IV. In contrast, Subclusters II and IV were composed exclusively of *U. brizantha* accessions, comprising 86 and 27 individuals, respectively, which collectively highlight the intra-specific diversity within this species. Subcluster III was a more heterogeneous group, containing accessions from all three species: *U. decumbens* (20%), *U. ruziziensis* (14%), and *U. brizantha* (8%). Similarly, Subcluster V represented a combination of species from *U. brizantha* (15%) and *U. decumbens* (11%). The admixture group, encompassing 89 accessions in total, including 30% *U. brizantha*, 20% *U. decumbens*, and the single Mulato II hybrid. These patterns are likely driven by evolutionary processes, historical polyploidization events that generated multiple ploidy levels within species, differences in reproductive mode (apomictic, facultative apomictic, or sexual), and ecological adaptation (Ryan et al., 2025; Higgins et al., 2022; Tomaszewska et al., 2023; Pessoa-Filho et al., 2017). Hybridization and gene flow among populations, potentially facilitated by domestication and cultivation practices, are also plausible contributing factors, particularly in the key forage species *U. brizantha* and *U. decumbens* (Tomaszewska et al., 2023; Higgins et al., 2022). Geographic patterns further reinforce the evolutionary and breeding history of Clade IV species. The majority of accessions originated from Eastern African countries, particularly Ethiopia, Kenya, Burundi, and Rwanda. *U. brizantha* accessions were predominantly from Ethiopia, followed by Kenya and Burundi; *U. decumbens* accessions were mainly from Rwanda and Kenya, with additional contributions from Burundi; and *U. ruziziensis* accessions were primarily from Burundi, followed by Rwanda and Kenya. This geographic structuring is consistent with the domestication and dissemination history of these species, reflecting both local adaptation and regional selection for forage productivity (Triviño et al., 2017; Worthington et al., 2021). Although most accessions analysed in this study were genebank collections that have not undergone formal breeding, a few improved cultivars, including the commercial cultivar Basilisk and the interspecific hybrid Mulato II, were included. Consequently, selective breeding may have contributed to the intricate population structure and widespread admixture observed in Clade IV, thereby enriching the reservoir of alleles associated with traits such as biomass yield, stress tolerance, and forage nutritional quality. The *Urochloa* breeding programs in different countries have effectively leveraged the extensive genetic diversity within *Urochloa* species (Mushagalusa et al., 2025; Worku et al., 2022; Ferreira et al., 2021; Leonardo et al., 2020; Jungmann et al., 2010). In particular, the programs at the Alliance of Bioversity–CIAT and EMBRAPA (Brazilian Agricultural Research Corporation) have made substantial progress in developing improved hybrids with enhanced pest resistance, superior nutritional quality, and broad adoption, contributing to significant economic gains and positive environmental impacts (Ferreira et al., 2021; Jank et al., 2014). Collectively, the results underscore the value of Clade IV species as a foundation for tropical forage improvement, emphasizing the importance of integrating natural and breeding-derived diversity to sustain productivity and resilience in livestock production systems under changing climatic conditions.

Overall, the clear separation of these four clades reflects the complex evolutionary history of the *Urochloa genus*, shaped by shared ancestry, ecological adaptation, polyploidization, and domestication processes. The cross-species markers identified in this study thus provide a valuable genomic framework for exploring genetic diversity, guiding breeding strategies, and conserving genetic resources across cultivated and wild *Urochloa* species.

### Population differentiation within *U. brizantha* accessions and progeny plants

*U. brizantha* represented the largest proportion of the population, with 253 accessions accounting for 46% of those analysed in this study. These accessions were grouped into three major clusters and five sub-clusters with an additional admixed group, consistent with previous findings (Higgins et al., 2022; Vigna et al., 2016). The pronounced population structure likely reflects the species’ broad geographic adaptation, variation in ploidy levels, and differences in reproductive mode (Higgins et al., 2022; Triviño et al., 2017), offering valuable opportunities for selecting genetically diverse parental genotypes. We observed an interesting association between trait variation and the genetic clusters identified through diversity analysis, suggesting a strong genetic basis underlying key morphological and phenological traits in the population. The differentiation among clusters for traits such as days to flowering, leaf length, plant height, and number of spikelet rows indicates that genetic structure may have influenced phenotypic divergence, potentially reflecting adaptation to different environments or parental line contributions (Ferreira et al., 2021). The genetic clusters in combination with the identified subset can serve as a reference set for distribution and evaluation across different agro-ecological zones and production systems. The subset accessions capture both the genetic diversity and geographical origins of the collection conserved in the ILRI Genebank. The genome-wide association study (GWAS) further supported the above findings by identifying consistent marker-trait associations, particularly the significant associations on chromosomes 1, 2, and 4 for leaf sheath hair, which explained 7–24% of the phenotypic variation. From a functional standpoint, leaf sheath hairiness can serve adaptive and agronomic roles, including defence against herbivory, tolerance to drought or heat stress, and microclimate regulation near the leaf surface (Saade et al., 2017).

The analysis of progeny plants derived from *U. brizantha* accessions revealed genetic uniformity among progenies from the same seed parent, as indicated by both Identity by Descent (IBD) analysis and Nei’s genetic distance estimates. The absence of detectable genetic variation within progeny groups, could be due to the highly restricted mode of reproduction, either apomictic or self-pollination, in *U. brizantha* (do Valle and Savidan 1996) both of which limit genetic recombination and result in genetically homogeneous offspring (Hojsgaard and Hörandl, 2019). While apomixis facilitates the propagation of elite genotypes without segregation, it also constrains the generation of novel genetic variation, limiting the potential for hybridization-based improvement (Ferreira et al., 2021; do Valle and Savidan 1996). Consequently, understanding the balance between apomictic and sexual reproduction in *U. brizantha* populations remains essential for designing effective breeding and germplasm enhancement strategies.

## Conclusions

This study provides a comprehensive assessment of genetic diversity, population structure, and evolutionary relationships across 17 *Urochloa* species using cross-species markers generated using the GBS DArTSeq platform. The accessions were clearly separated into four major clades, distinguishing the brizantha complex (*U. brizantha, U. decumbens,* and *U. ruziziensis*) composed of the major cultivated species from the rest of the species mainly composed of wild relatives, while revealing substantial intra- and inter-specific variation.

The wild species exhibited strong genetic differentiation with limited admixture, often reflecting geographic origin, highlighting their value as reservoirs of adaptive traits such as drought tolerance, waterlogging resilience, and disease resistance. In contrast, the cultivated species displayed extensive admixture and subpopulation structuring, including the interspecific hybrid, consistent with historical shared ancestry, polyploidization events and domestication processes.

The observed patterns of genetic diversity and population structure have significant implications for conservation, germplasm management, and breeding strategies. Wild species provide unique alleles for stress adaptation and resilience, while the diverse cultivated gene pool offers opportunities to enhance forage productivity and nutritional quality through targeted hybridization and selection. Overall, this study lays a foundation for strategic utilization of *Urochloa* genetic resources maintained in the ILRI Forage Genebank to support climate-smart tropical forage systems and sustainable livestock production.

## Acknowledgements

The authors gratefully acknowledge the ILRI Forage Genebank for providing the germplasm used in this study. The authors also thank Mr. Yonas Sime for preparing the map showing the germplasm collection sites.

## Author Contributions

C.S.J., M.S.M., and A.T.N. designed and supervised the project and the manuscript writing; M.S.M. and S.D.L. analyzed the data and wrote the manuscript; Y.A. and H.M.T. collected the phenotype data; S.D.L. and Y.A. collected leaf samples and extracted DNA; H.N. involved in the supervision of the data analysis and manuscript writing. All authors made a significant contribution to the development of this manuscript and approve it for publication.

